# Highly efficient scarless knock-in of reporter genes into human and mouse pluripotent stem cells via transient antibiotic selection

**DOI:** 10.1101/374728

**Authors:** Valentin M. Sluch, Xitiz Chamling, Claire Wenger, Yukan Duan, Dennis S. Rice, Donald J. Zack

**Affiliations:** Department of Ophthalmology, Novartis Institutes for BioMedical Research, Cambridge, Massachusetts, USA; Department of Ophthalmology, Wilmer Eye Institute, Johns Hopkins University School of Medicine, Baltimore, Maryland, USA; McKusick-Nathans Institute of Genetic Medicine, Johns Hopkins University School of Medicine Baltimore, Maryland, USA; Department of Molecular Biology and Genetics, The Solomon H. Snyder Department of Neuroscience, Institute of Genetic Medicine, Johns Hopkins University School of Medicine, Baltimore, Maryland, USA

**Keywords:** CRISPR-Cas9, pluripotent stem cells, homology directed repair, knock-in, puromycin selection

## Abstract

Pluripotent stem cells (PSCs) edited with genetic reporters are useful tools for differentiation analysis and for isolation of specific cell populations for study. Reporter integration into the genome is now commonly achieved by targeted DNA nuclease-enhanced homology directed repair (HDR). However, human PSCs are known to have a low frequency of gene knock-in (KI) by HDR, making reporter line generation an arduous process. Here, we report a methodology for scarless KI of large fluorescent reporter genes into PSCs by transient selection with puromycin or zeocin. With this method, we can perform targeted KI of a single reporter gene with up to 65% efficiency, as well as simultaneous KI of two reporter genes into different loci with up to 11% efficiency. Additionally, we demonstrate that this method also works in mouse PSCs.

## Introduction

Pluripotent stem cells (PSCs) represent a powerful tool for disease modeling as well as *ex vivo* developmental and mechanistic studies, and they are also currently being used in clinical trials for regenerative medicine(1-7). Due to advances in genome editing technologies, such as zinc finger nucleases, transcription activator-like effector nucleases, and clustered regularly interspaced short palindromic repeats with the associated Cas9 protein (CRISPR-Cas9)(8), it has become possible to routinely perform genome editing in PSCs. Targeted mutations for disease modeling(9) or knock-in reporter genes(10-12) to aide in high throughput screening(13) or to mark cells for isolation have been introduced into PSCs via genome editing. Nevertheless, genome editing of human PSCs, including both induced pluripotent stem cells (hiPSCs) and embryonic stem cells (hESCs), remains challenging due to the low efficiency of homology directed repair (HDR)-based knock-in (KI) in PSCs, with frequencies reported to be generally around 1%(14, 15). A number of groups have attempted to overcome this low efficiency by enriching for transfected stem cells via fluorescence activated cell sorting (FACS)(16, 17), enhancing HDR frequencies via small molecule treatment(18) or synchronizing the cell cycle (19), or utilizing non-homologous end joining (NHEJ) pathways for KI(20). These efforts have indeed helped to improve the efficiency of single base pair edits. For example, by using FACS-based enrichment of transfected cells, or permanent integration of Cas9 into the genome, single base pair edits have been reported with 11% and 34% efficiency, respectively(16, 21). However, FACS instruments for live-cell sorting are not available to every lab, and the genomic integration of Cas9 into cells may have long-term detrimental effects due to potential Cas9 toxicity(22) and the risk of non-specific DNA cleavage(23). Moreover, efficient, targeted KI of larger DNA elements, such as fluorescent reporter genes, has not been demonstrated in these studies(16, 21).

The co-introduction of a gene conferring antibiotic resistance with the target KI gene allows for selection of the KI event, and has been suggested to be a more efficacious strategy to facilitate integration of the target into the genome(24, 25). This strategy can be highly efficient, with reported KI frequencies of 1.5 to 94%, although with significant variability across cell lines and target loci. Despite the increased KI efficiency associated with this method, a potential concern is that the integrated antibiotic resistance gene remains as a permanent genetic “scar” that may unintentionally affect nearby gene expression(26). Although it is sometimes possible to remove this additional exogenous DNA, e.g. using Cre-lox or FLP-FRT recombinase technologies(24) or transposases(27), this treatment prolongs the required experimental time and effort, and adds additional DNA manipulation into the experimental workflow. Here, we report an improved method for “scarless” KI of large constructs, such as fluorescent reporters, by simply enriching for cells transfected with the Cas9-P2A-Puro plasmid via transient puromycin selection. We show that a transient, 24-hour antibiotic selection dramatically increases the KI frequency of large inserts, such as eGFP, by over 33-fold and with an efficiency approaching ~65%. We validate the efficacy of this method across multiple targets, inserts, and stem cell lines, and show that simultaneous KI of two reporter genes is also achievable. Furthermore, this method improves KI efficiency in mouse embryonic stem cells (mESCs), up to ~60%, and it is not limited to the use of puromycin, as use of zeocin yields a similarly high efficacy.

## Results

### Transient puromycin selection facilitates efficient KI of eGFP into hESCs

PSCs are traditionally transfected using electroporation-related methodologies. In an effort to develop a simpler, more cost-effective, and higher throughput method, we utilized DNA-In Stem, a lipofection reagent (GST-2130, MTI-GlobalStem, Thermo Fisher Scientific). With DNA-In Stem, we achieved 25-30% transfection of large plasmids such as lenti-CMV-eGFP (9 kb) and Cas9-P2A-mKate2 (Fig 1). In order to further enrich for Cas9-transfected cells and to improve our chances of isolating HDR-corrected clones, we transfected H9 hESCs with a Cas9-P2A-Puro+U6-gRNA plasmid and selected the cells with puromycin after transfection. We established 0.6 μg/mL as an optimal dose of puromycin for H9 hESCs to ensure that all untransfected cells were dead after a 24-hour treatment. To test if puromycin-based selection of the Cas9-transfected hESCs improves the KI efficiency of large DNA sequences, we performed a targeted KI of an *H2B-eGFP* sequence into the C-terminal end of the housekeeping gene encoding TATA-box binding protein (*TBP*) (Fig 2A). In this setup, the cells that undergo HDR-based KI of *H2B-eGFP* express a nuclear localized eGFP (Fig 2B). Forty hours after transfection, cells were treated with puromycin for 24 hours and then allowed to recover and expand for 5-7 days. Successful KI was then assessed using fluorescent microscopy and quantified using flow cytometry analysis. With puromycin selection, we achieved a remarkably high KI frequency of ~46.6% compared to ~1.4% without selection (Fig 2C, D). For simplicity, we refer to all subsequent KI sequences by their fluorescent gene name only, and their additional cell localization details can be found in the methods section.

**Fig 1.**
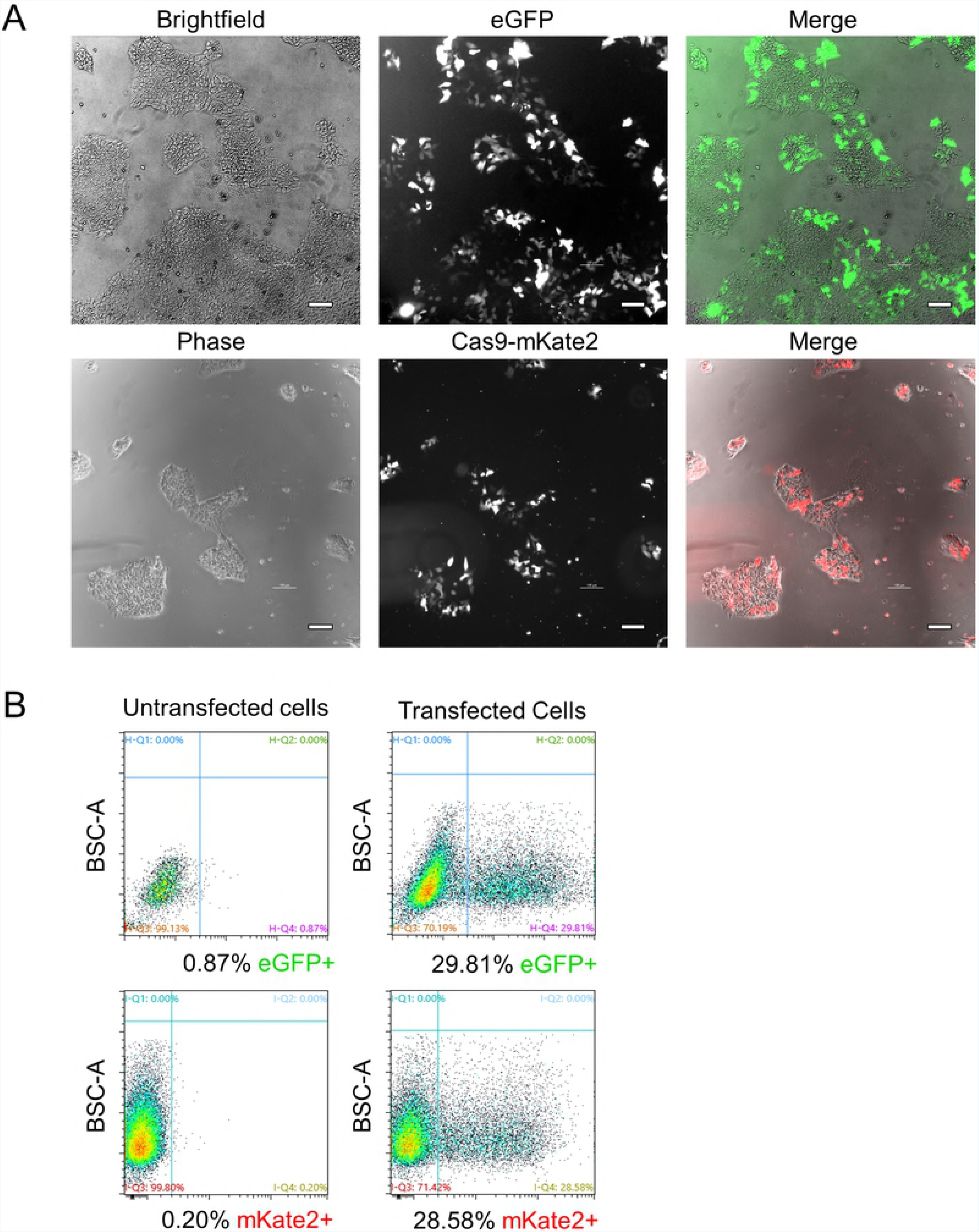
Transfection efficiency of H9 hESCs using lipofection. (A) Phase/brightfield and fluorescence microscopy images of eGFP or Cas9-mKate2 plasmid transfected H9 hESCs are shown. Scale bar = 100 μm. (B) Flow cytometry assessment of H9 hESCs transfection. Untransfected cells were used to set the negative gates. BSC-A = back scatter area. Cells were analyzed at 40 hours post transfection in both A and B.

**Fig 2.**
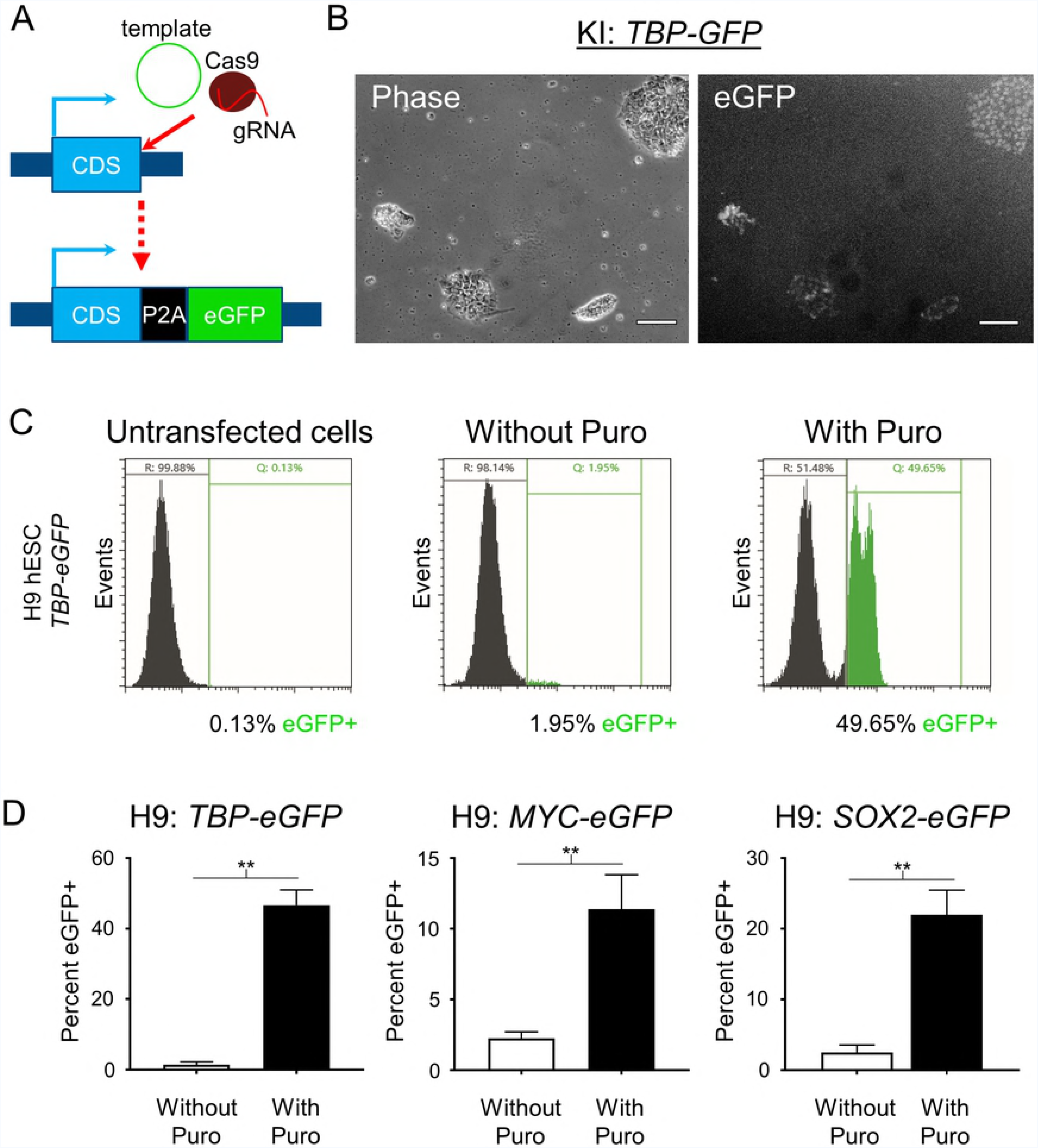
Transient puromycin selection increases reporter KI efficiency. (A) Schematic of gene reporter design created by CRISPR editing. (B) Phase and fluorescence microscopy images of *TBP-P2A-eGFP* KI into H9 hESCs after puromycin selection. Scale bar = 100 μm. (C) Representative images of flow cytometry assessment of *TBP-P2A-eGFP* KI into H9 hESCs with and without transient puromycin selection. Untransfected cells were used to set the gates for reporter negative cells. Two peaks of eGFP+ cells can be observed in the puromycin treated group suggesting homozygous and heterozygous KI. (D) Flow cytometry analysis of *TBP-P2A-eGFP*, *MYC-P2A-eGFP*, and *SOX2-P2A-eGFP* KI into H9 hESCs with and without transient puromycin selection. For *TBP* n = 2 for both groups, for *MYC* n = 2 for replicates without puromycin and n = 5 for replicates with puromycin treatment, for *SOX2* n = 3 for replicates without puromycin and n = 4 for replicates with puromycin treatment. n = biological replicates. *p* values: *TBP* = 0.0047, *MYC* = 0.0041, *SOX2* = 0.0057. ** = *p*<.01. Unpaired two tailed t-test was used.

### Transient puromycin selection increases KI efficiency with multiple genes and cell lines

To test the generality of the increased HDR efficiency that we noted with *TBP* in H9 hESCs, we tested additional P2A-eGFP KI gene reporters (MYC proto-oncogene, bHLH transcription factor (*MYC*) and SRY-box 2 (*SOX2*)). Although the efficiency of KI for these genes was lower than *TBP* in H9 hESCs, puromycin treatment successfully increased the KI frequency from 2.3% to 11.4% and from 2.5% to 22.0% for *MYC* and *SOX2*, respectively (Fig 2D). We also confirmed that transient puromycin selection increases KI efficiency in other hESC and hiPSC lines: H7, IMR90-4 and EP1(28, 29). Although some expected variability in KI efficiency for the different lines was noted, transient puromycin selection yielded a 5 to 24-fold increase in KI frequencies for all hESC and hiPSC lines tested (Fig 3). Then to test KI for a gene that is not expressed in undifferentiated stem cells, since chromatin state associated with transcriptional activity could potentially affect HDR efficiency, we transfected EP1 hiPSCs with the *BRN3B* (*POU4F2*)*-P2A-tdTomato-P2A-Thy1.2* construct that we previously used to make retinal ganglion cell (RGC) reporter lines in H7 and H9 hESCs(11). Following transfection and transient puromycin selection, we plated the surviving cells as single cells at a low density for clonal derivation. PCR-based genotyping of 65 individual clones showed a markedly increased KI efficiency of 64.6% (Fig 4), compared to our initial frequency of 1.4% using traditional methodology with this same reporter construct(11). Karyotyping and PCR-based off-target analysis of the EP1-derived *BRN3B* reporter line showed that the puromycin treatment did not cause chromosomal abnormalities or off-target editing (S1 Fig)(12). Additionally, we differentiated the EP1 reporter line to RGCs per our prior protocol(11), and observed no differences in our ability to derive RGCs from this reporter line generated using transient puromycin selection.

**Fig 3.**
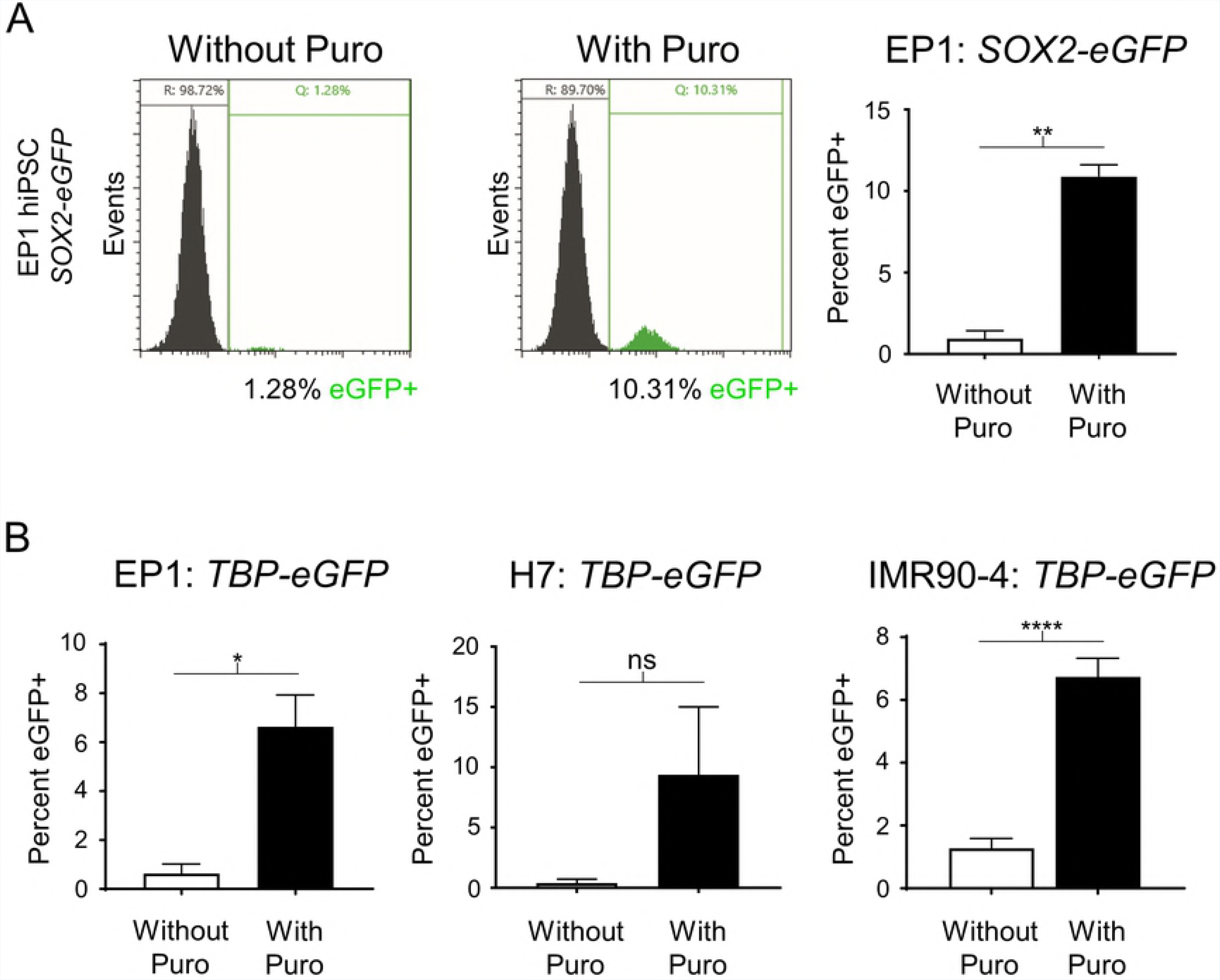
Demonstration of increased reporter KI efficiency in multiple human PSC lines after transient puromycin treatment. (A) Representative images of flow cytometry assessment of *SOX2-P2A-eGFP* KI into EP1 hiPSCs with and without transient puromycin selection followed by flow cytometry analysis. Untransfected cells were used to set the gates for reporter negative cells. n = 2 for both groups. *p* value = 0.0039. (B) Flow cytometry analysis of *TBP-P2A-eGFP* KI into EP1 hiPSC, H7 hESC, and IMR90-4 hiPSC lines, respectively, with and without transient puromycin selection. For EP1 and H7, n = 2 for both groups, for IMR90-4 n = 3 for replicates without puromycin and n = 4 for replicates with puromycin treatment. n = biological replicates. *p* values: EP1 = 0.0246, H7 = 0.1532, IMR90-4 = <0.0001. * = p<.05, ** = p<.01, **** = p<.0001. ns = not significant. Unpaired two tailed t-test was used.

**Fig 4.**
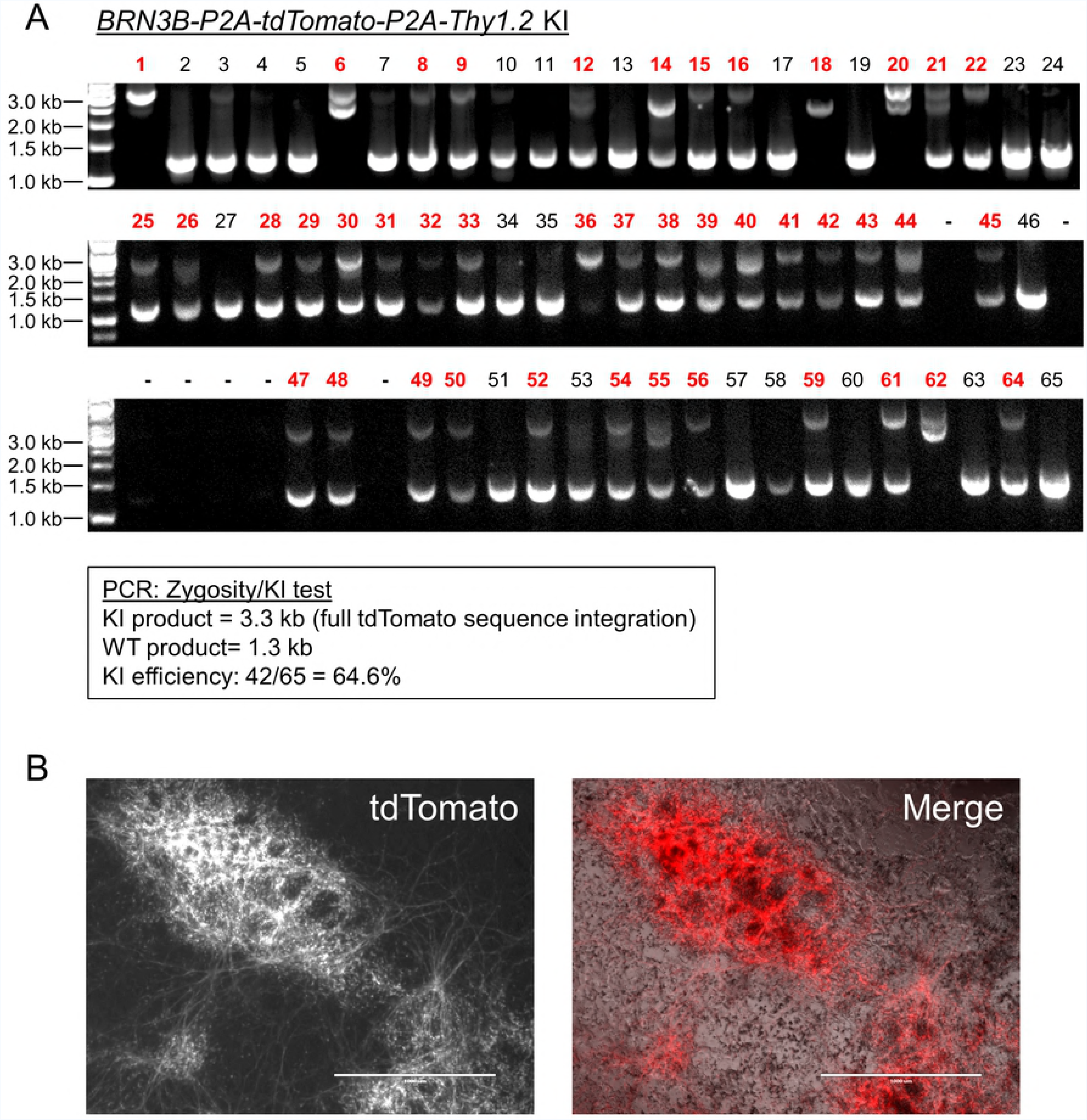
Generation of an RGC reporter line in EP1 hiPSC background using transient puromycin treatment. (A) PCR zygosity test for KI at the targeted *BRN3B* locus. Primers spanning the integration region were used to amplify genomic DNA from randomly picked colonies derived from plating single cells. Homozygous insertion of the KI cassette is indicated by a single band at 3.3 kilobase pairs (kb). KI negative clones generate a band of 1.3 kb. Clones producing both bands were scored as heterozygous KI. WT = wildtype. For some of the clones (e.g. lane 6), the KI product is split into two parts due to an incorporation of only one monomer of the tdTomato sequence. (B) Fluorescence and phase microscopy of a differentiated EP1 hiPSC RGC reporter line generated using transient puromycin selection. Cells were imaged on day 29 of differentiation. Scale bar = 1000 μm.

### Transient puromycin selection allows for the creation of dual reporter KI lines

Based on the ability of puromycin transient selection to promote generation of single reporter KI lines, we evaluated its ability to promote simultaneous KI of two reporters in two different loci. For this experiment, we targeted a KI of *TBP-P2A-eGFP* and *MYC-P2A-tdTomato* or *TBP-P2A-tdTomato* and *SOX2-P2A-eGFP* into H9 hESCs and analyzed the result by flow cytometry. With puromycin selection, KI frequencies were ~9.5% and 11.3% for each combination, respectively (Fig 5). Notably, most of our KI cells were double positive, supporting the previously reported observation that HDR occurrence at one locus is associated with an increased frequency of HDR at other loci(30, 31).

**Fig 5.**
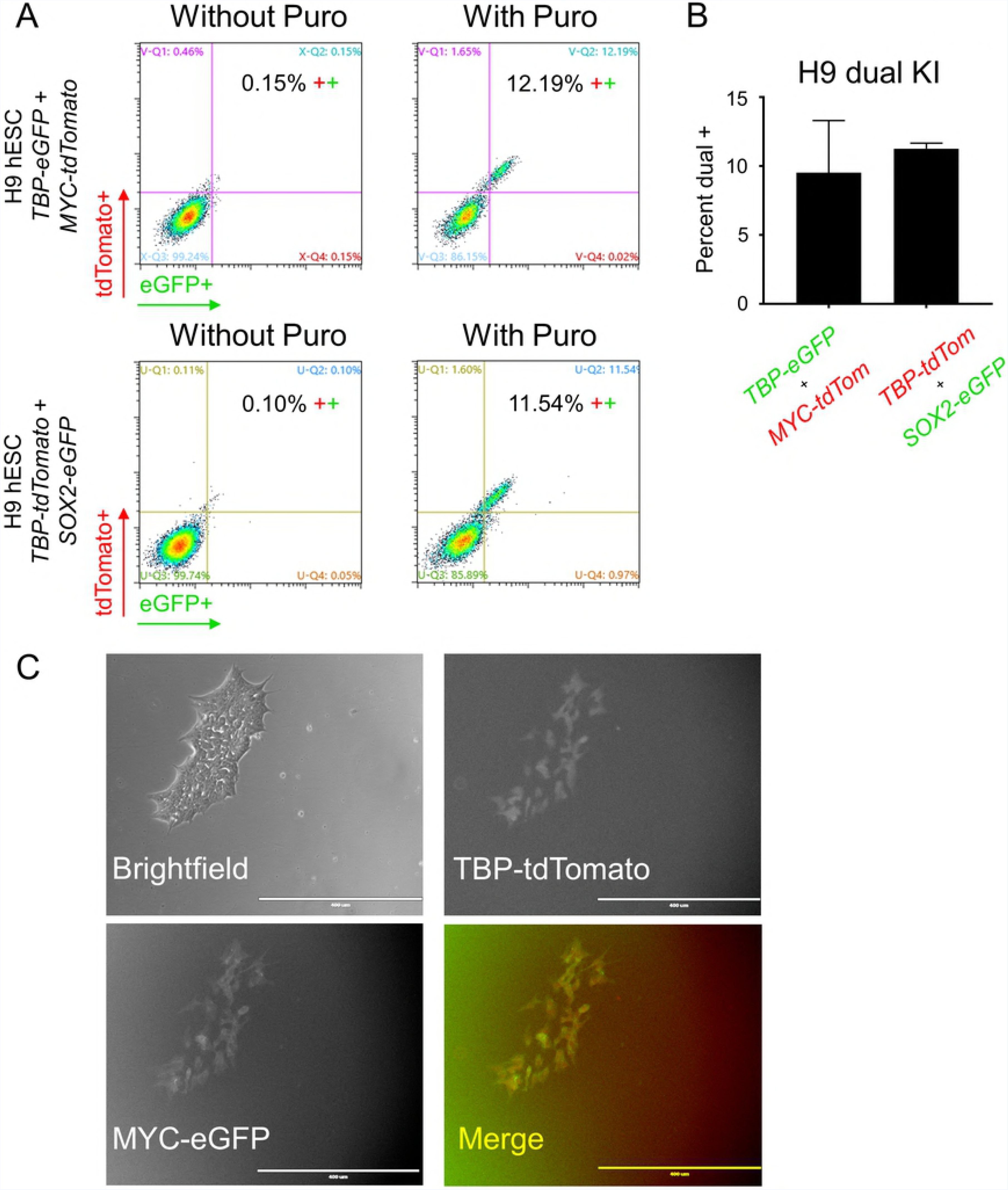
Dual KI of *TBP* and *MYC* or *TBP* and *SOX2* fluorescent reporters into H9 hESCs. (A) Representative images of flow cytometry assessment of dual KI of *TBP-P2A-eGFP* and *MYC-P2A-tdTomato* or *TBP-P2A-tdTomato* and *SOX2-P2A-eGFP* into H9 hESCs with and without transient puromycin selection. (B) Flow cytometry analysis of the dual KI described in A. n = 2 for both dual KI combinations. n = biological replicates. (C) Fluorescence and brightfield microscopy of H9 hESCs positive for both reporter genes. Scale bar = 400 μm.

### Transient puromycin selection improves KI in mESCs

Given that all the above-described experiments were performed with human PSCs, we wanted to test if transient puromycin selection would also promote HDR in mouse stem cells. After establishing an appropriate puromycin dose for the commonly used mouse embryonic stem cell (mESC) line E14TG2a(32), we transfected these cells via lipofection with CRISPR plasmids and the corresponding donor templates for *AdipoR1-eGFP*, *Mitf-P2A-tagRFP*, and *Six6-P2A-GFP*, targeting the genes for the adiponectin receptor 1, melanogenesis associated transcription factor, and sine oculis-related homeobox 6, respectively. Due to their faster growth rate, we treated the mESCs with puromycin at 24 hours post transfection rather than the 40 hours that was used for human PSCs. Following recovery, the surviving populations were plated as single cells to derive independent colonies and 24 colonies were PCR-screened for KI insertion. We found KI frequencies of 45.8%, 33.3%, and 58.3% for targeting *AdipoR1*, *Mitf*, and *Six6*, respectively (Fig 6). To check whether puromycin treatment had abrogated the ability of these cells to faithfully differentiate, we tested one of the homozygous *Six6-P2A-eGFP* lines in optic cup retinal differentiation(33). *Six6* is an eye field transcription factor that is highly expressed in the optic vesicle(34, 35) during retinal development. After 8 days of differentiation, we observed eGFP positive vesicles emerge from the main aggregate signifying successful retinal differentiation.

**Fig 6.**
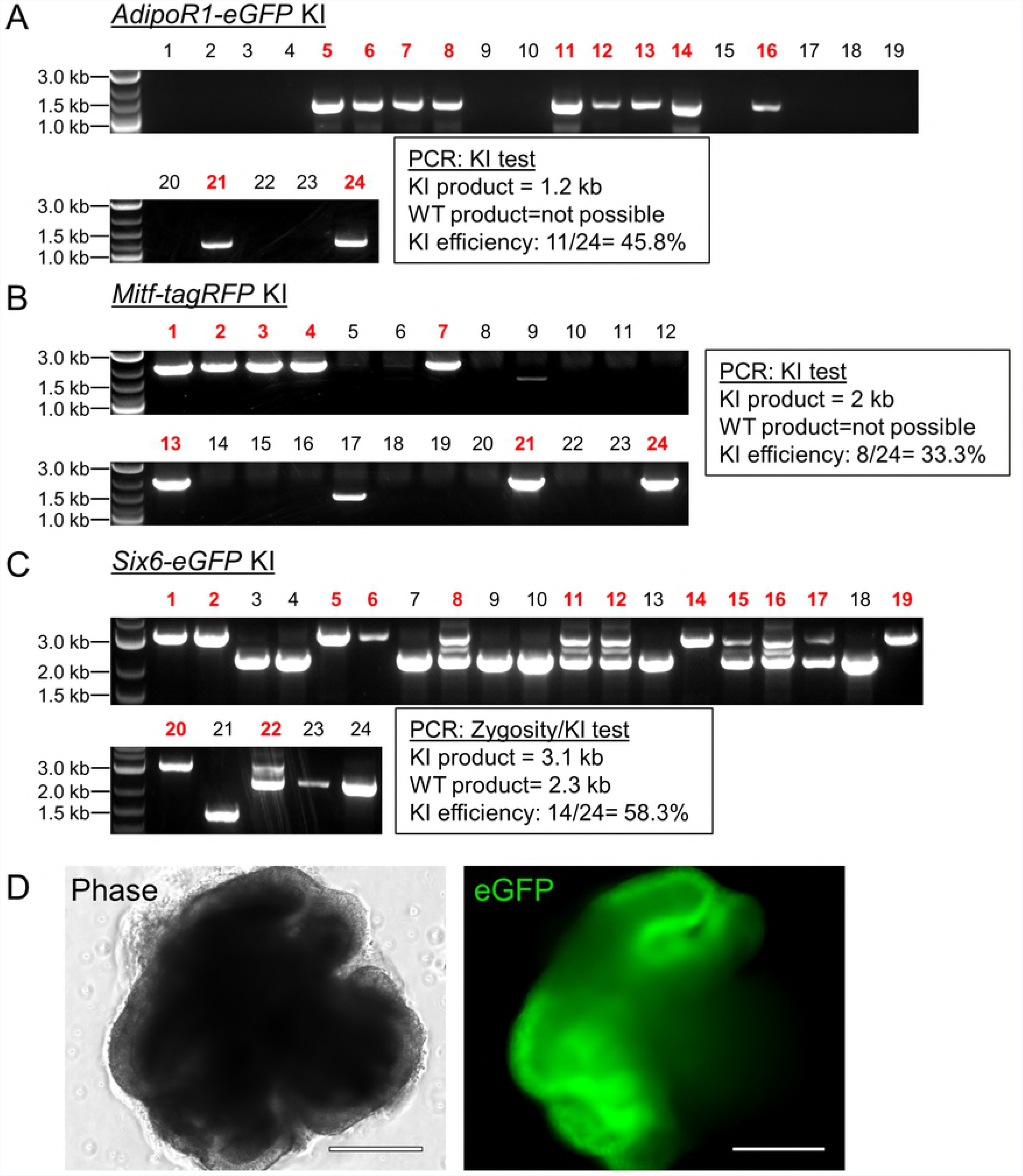
Transient puromycin selection results in high KI efficiency of fluorescence reporter genes into mESCs. (A, B, C) PCR tests for fluorescent reporter KI at the indicated loci in mESCs. For A and B, primers amplifying a region inside the KI gene and outside the donor plasmid template were used. For C, primers spanning the integration region were used to distinguish between homozygous and heterozygous clones. Expected amplicon sizes are shown. WT = wildtype. For KI assessment in B, lanes 9 and 17 were not counted as positive KI because the amplicon did not run at the predicted size. (D) Phase and fluorescence microscopy of a homozygous *Six6-P2A-eGFP* reporter KI line generated in C. mESCs were differentiated to optic vesicles for 8 days. Scale bar = 275 μm.

### Zeocin can replace puromycin for transient selection-based KI

Next, to test if the ability of transient puromycin selection to improve KI efficiency is simply due to enrichment of transfected cells or through some other activity specific to puromycin, we tested if treatment with another selective agent would increase KI efficiency to a comparable extent. We replaced the puromycin resistance sequence in the Cas9 plasmid with one for zeocin resistance (Cas9-P2A-Zeocin). Zeocin, which has a completely different mechanism of action (targeting DNA) than puromycin (inhibiting protein translation), also results in rapid death of untransfected stem cells(36). We repeated the KI experiments using Cas9-P2A-Zeocin and the *TBP-P2A-eGFP* donor plasmids. Transient 24 hour zeocin selection at 40 hours post transfection resulted in 13.7% and 9.2% KI efficiency for IMR90-4 hiPSCs and H7 hESCs, respectively (Fig 7), values that were actually slightly higher than our prior puromycin selection efforts targeting *TBP* in these cell lines.

**Fig 7.**
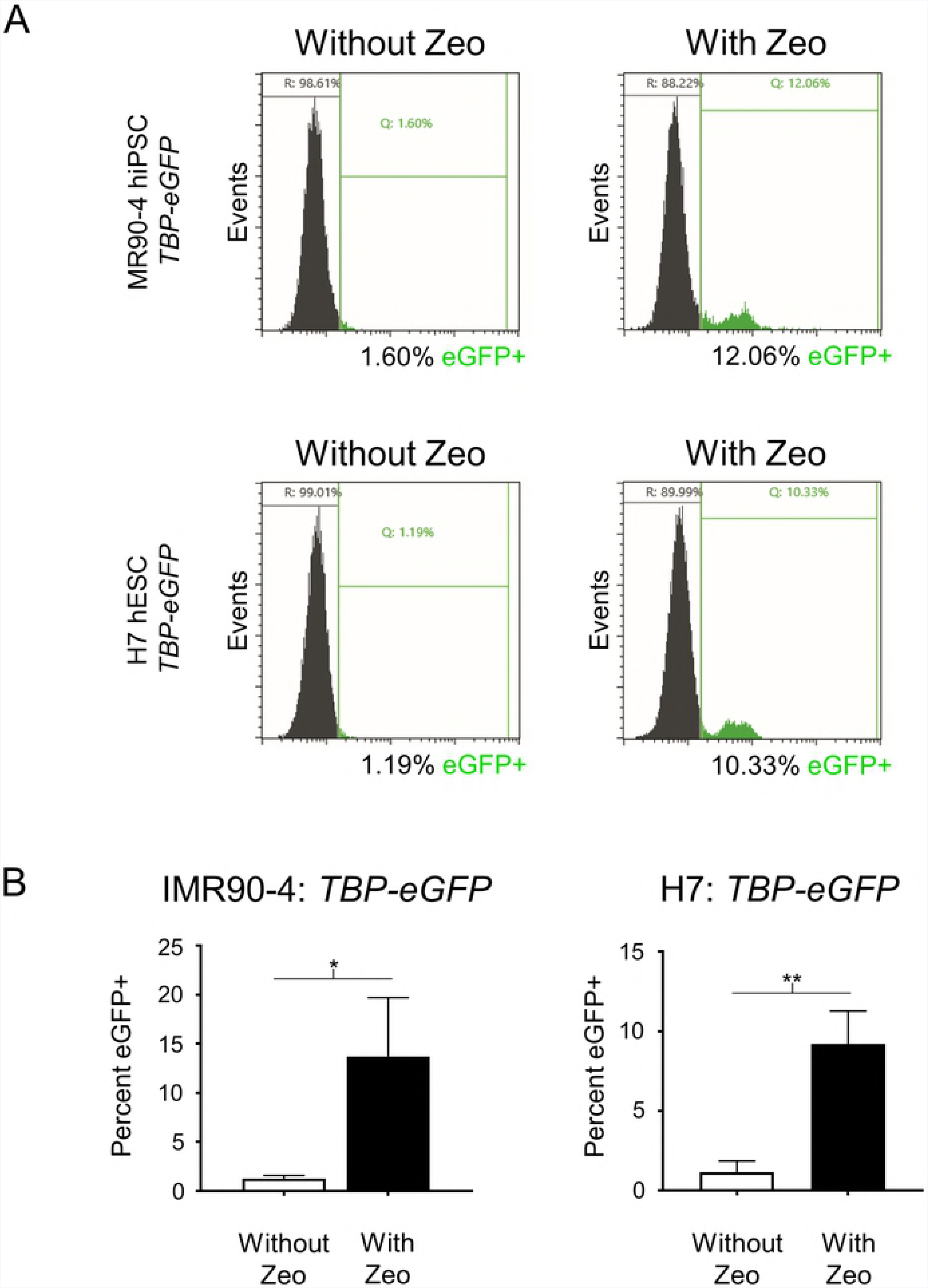
Zeocin replacement of puromycin results in similar KI efficiency of fluorescent reporter genes into human PSCs. (A) Representative images of flow cytometry assessment of *TBP-P2A-eGFP* KI into IMR90-4 hiPSCs and H7 hESCs with and without transient zeocin selection. (B) Flow cytometry analysis of *TBP-P2A-eGFP* KI in part A. For IMR90-4, n = 3 for replicates without zeocin and n = 4 for replicates with zeocin treatment, for H7 n = 2 for replicates without zeocin and n = 5 for replicates with zeocin treatment. n = biological replicates. *p* values: IMR90-4 = 0.0172, H7 = 0.0036; * = p<.05, ** = *p*<.01; Unpaired two tailed t-test was used.

## Discussion

Genome editing of pluripotent stem cells (PSCs) greatly enhances the utility of PSCs for studying biological processes, use in disease modeling(9), and reporter line generation for specific cell type isolation(11, 12). Here, we have demonstrated that transient selection with puromycin or zeocin can facilitate highly efficient scarless KI of relatively large reporter genes (over 2 kb in the case of *BRN3B-P2A-tdTomato-P2A-Thy1.2*) into both human and mouse PSCs. These KI cell lines differentiated normally, and did not show karyotypic abnormalities or display off-target effects. Moreover, since the antibiotic resistance gene is not integrated into the genome, these reporter-positive clones could be directly expanded for use in experiments without the need to undergo further manipulation to excise the resistance cassette.

We have successfully applied the transient selection method in four human PSC lines for KI at four different loci as well as one mESC line with KI at three other loci. Our observed KI efficiencies varied from 6.6 to 64.6% in human PSCs and from 33.3 to 58.3% in mESCs, presumably reflecting the high degree of gene editing variability that has also been observed by others (24, 25). Although mouse stem cells generally demonstrate higher KI frequencies than their human counterparts (e.g. a previous study had reported a KI efficiency of 15% for insertion of eGFP(37)), the KI frequencies we observed in mESCs were even higher than the ones achieved via small molecules(18) or positive KI selection(31). With the high KI efficiencies described in this manuscript, with both human and mouse PSCs, it becomes reasonable to isolate reporter-positive lines from as few as approximately twenty clones versus the standard practice of picking hundreds of clones. Moreover, we were able to generate dual KI reporter cell lines at ~10% efficiency. A dual KI reporter could be advantageous to mark different cell subtypes(11), label differentiation progression(38), or to increase stringency of selection in situations where a single gene signature is not enough to define a specific cell population. Importantly, based on our data and the previous report that successful HDR at one locus is associated with increased frequency of HDR at additional loci(30, 31), it should be possible to multi-plex reporter KIs to more than two loci. Additionally, with a high KI efficiency it may be advantageous to knock-out genes by knocking-in stop codon sequences at precise locations instead of relying on NHEJ to generate loss of function alleles by random mutagenesis.

Recently, Steyer and Bu et al.(39) reported that transient puromycin selection for 3 consecutive days increased the efficiency of single base pair KI in human PSCs to values between 14 to 44%. Notably, as KI efficiency tends to decrease with increasing insert size(40), their reported KI frequency for a 21 base pair insert was a little lower at 36%, and they did not demonstrate that their method could be used to KI larger constructs. In their method, human PSCs are electroporated and puromycin is added for 3 days starting at 24 hours post transfection along with a Rho kinase (ROCK) inhibitor. Our methodology utilizes lipofection and puromycin/zeocin that was added at ~40 hours post transfection and for only 24 hours. Although both our data and their work(39) suggest that puromycin treatment does not alter the karyotype of PSCs, or increase off-target effects, decreasing the exposure of the cells to puromycin is likely to reduce the possibility of undesired side-effects, as prolonged antibiotic exposure could increase the risk of random integration of the resistance gene into the genome.

We wondered if the observed KI efficiency was increased due to a more robust selection pressure that was enriching for the highest Cas9-P2A-Puro expressing cells or whether puromycin itself had an effect on HDR. To explore this hypothesis we tested whether zeocin, an antibiotic with a different mechanism of action from puromycin, would yield a comparable high KI frequency to that which we had observed with puromycin, and found that it did. This result argues for selection pressure being the main driver of improved KI efficiency. Similarly, Steyer and Bu et al. have reported that utilizing FACS to enrich for the highest Cas9-2A-eGFP-expressing cells could increase KI frequencies to levels comparable to those achieved by transient puromycin selection, further supporting the selection pressure hypothesis.

In addition to our work and Steyer and Bu et al., other methods have also been recently introduced to increase KI efficiency in PSCs. Guo et al.(41) have reported that a cold-shock of hiPSCs achieved by decreasing the culture temperature to 32 °C for 24–48?hours following transfection increased KI of small base pair inserts to 20–40%. However, this cold-shock appeared to have a negative effect on NANOG expression, a potential sign of decreased pluripotency. Notably, Zhang and Li et al.(42) introduced a method that could improve KI efficiency for large constructs to 15-30% in hiPSCs. However, to generate this effect their experimental workflow requires cell cycle synchronization via a co-transfection with a *CCND1* plasmid and Nocodazole treatment in combination with a pre-designed donor vector containing gRNA sites that enable donor template linearization inside the cell. Importantly, the effects of these treatments on the karyotype, off-target mutations, and downstream differentiation were not addressed in either study. It is possible that the combinatorial effect of cold-shocking or cell cycle synchronization would also translate to an improved KI efficiency when combined with our transient antibiotic selection. However, in comparison to these previous reports, the method described here is already highly efficient, technically simpler, requires less manipulation, and can be adapted by any lab with a basic cell culture facility.

## Methods

### Plasmid design

We used the following gRNAs for targeting their respective loci (5’-3’, PAM sequence in bold)

*BRN3B* – GCCAAGAGTCTTCTAAATGC **CGG**

*TBP* – GATTCAGGAAGACGACGTAA **TGG**

*MYC* – AACACAAACTTGAACAGCTA **CGG**

*SOX2* – GGCCCTCACATGTGTGAGAG **GGG**

*AdipoR1* - GGCTCAGAGAAGGGAGTCGT **CGG**

*Mitf* - GAGCTTCAAAAACAAGCAGC **CGG**

*Six6* - GATGTCGCACTCACTGTCGC **TGG**

The gRNA sequences were based on our prior publications(11, 12) or designed using CRISPRdirect(43) (https://crispr.dbcls.jp) and cloned into an all-in-one (CMV-Cas9 + U6-gRNA) Cas9-P2A-Puro or Cas9-P2A-Zeo plasmid (modified from Addgene #62988 plasmid to replace T2A with a P2A sequence).

New donor plasmids were generated via PCR amplification of human or mouse genomic DNA of approximately 1000 base pairs on each side of the gRNA target site. This homology template was then cloned into Zero Blunt vectors by TOPO cloning (Thermo Fisher Scientific, https://www.thermofisher.com). The reporter gene was then introduced into these donor vectors using Gibson assembly (NEB, https://www.neb.com). By design all gRNA binding genomic sequences are destroyed by integration of the reporter into the genome to prevent further CRISPR cutting.

The following donor vectors were used in this study:

BRN3B-P2A-tdTomato-P2A-Thy1.2

MYC-P2A-tdTomato

MYC-P2A-myr-eGFP (myr is a myristoylation sequence for membrane localization)

SOX2-P2A-eGFP-nls (nls is a nuclear localization signal from SV40)

TBP-P2A-H2B-eGFP (H2B is a nuclear localization signal from Histone H2B)

TBP-P2A-tdTomato

AdipoR1 fusion – eGFP (a truncated P2A was used as a fusion linker)

Mitf-P2A-tagRFP

Six6-P2A-eGFP-nls

### PCR KI Screening

After genome editing and puromycin selection, stem cells were expanded and passaged as single cells. Formed colonies were individually picked and screened for reporter integration by PCR using the following forward and reverse primers (5’-3’):

#### BRN3B

Primers amplify a region of human genomic DNA spanning the KI integration site. WT DNA produces a 1.3 kb region and KI reporter DNA produces a 3.3 kb region.

forward: GGAGAAGCTGGACCTGAAGAAAAACGTG
reverse: CCTTGGTGAAATCTAAAATCTGAAGGG

#### AdipoR1

Primers amplify a 1.2 kb region of mouse genomic DNA outside the homology plasmid and a region of fusion-tag+eGFP

forward: CTTTCTATGATCTTAATGGGAATCTACTCTTCTGGCTTTG
reverse: TCGCCCTTGCTCACCATAGGGCCGGGGTTCTCCTCCACGTCG

#### Mitf

Primers amplify a 2 kb region of mouse genomic DNA outside the homology plasmid and a region of tagRFP

forward: GACCCTGAATACAGCTCTTTTGTGTAGGCATCTC
reverse: CAGGCTCGCTATCAATTAAGTTTGTGCCCCAGTTTGCTAGGGAGG

#### Six6

Primers amplify a region of mouse genomic DNA spanning the KI integration site. WT DNA produces a 2.3 kb region and KI reporter DNA produces a 3.1 kb region.

forward: CGGGAGGAGGCATTCTTGGCCCTTA
reverse: GCCTGCATACTGTCTCCTATCTTAGTATTTCTCCTGGTG

### Cell culture

Human PSCs (EP1(28, 29) or H7, H9, IMR90-4, WiCell, https://www.wicell.org) were maintained by clonal propagation in mTeSR1 media (Stemcell Technologies, https://www.stemcell.com) on growth factor-reduced Matrigel coated plates (354230, Corning, https://ecatalog.corning.com) at 10% CO2/5% O2. Cells were passaged by dissociation with Accutase (A6964, Sigma-Aldrich, https://www.sigmaaldrich.com) or TrypLE Express (12605010, Thermo Fisher Scientific). mTeSR1 media containing 5 mM blebbistatin (Sigma-Aldrich) was used for maintenance of single cells.

Mouse ESCs (ES-E14TG2a, ATCC CRL-1821, https://www.atcc.org) were maintained on Matrigel coated plates in 2i media - [50:50 mix of Neurobasal (21103049, Thermo Fisher Scientific):DMEM/F12 (10565042, Thermo Fisher Scientific) with 0.5% N2 (17502001, Thermo Fisher Scientific), 1% B27 (17504044, Thermo Fisher Scientific), 1% Anti-anti (15240062, Thermo Fisher Scientific), 0.5% Glutamax (35050061, Thermo Fisher Scientific), 0.1mM β- mercaptoethanol, 1000U/mL LIF (ESG1107, Millipore, http://www.emdmillipore.com), 1 μM PD0325901 (Selleck Chemicals, http://www.selleckchem.com), 3 μM CHIR99021 (Selleck Chemicals).

Puromycin/zeocin killing doses were established by plating single cells on Matrigel coated plates (with ROCK inhibitor for human stem cells). The following day cells were treated with culture media containing increasing doses of puromycin/zeocin. After 24 hours, cells were viewed under the microscope and the lowest dose resulting in ~100% cell death was selected for future KI experiments. For E14TG2a mESCs a puromycin dose of 3 μg/mL was used. For H7 and H9, 0.6 μg/mL of puromycin or 25 μg/mL of zeocin; and for IMR90-4 and EP1 0.9 μg/mL of puromycin or 40 μg/mL of zeocin was used. Since the optimal dose can vary, we recommend performing dose response experiments for each cell type.

### Transfection and transient selection KI

#### Human stem cells

Human PSCs were plated at 50K per well of a 24 well plate in mTeSR1 media with ROCK inhibitor, blebbistatin. The following day, (for one well) 0.35 μg of all-in-one Cas9 plasmid and 0.75 μg of donor plasmid were combined with Opti-MEM (31985062, Thermo Fisher Scientific) to a total volume of 48 μL. For transfection tests, Cas9-P2A-mKate2 or lenti-CMV-eGFP (9 kb) plasmids were used. Cas9-P2A-mKate2 was modified from PX458 (Addgene #48138) by replacing GFP with mKate2. Transfection mix was prepared by adding 2 μL of DNA-In Stem (GST-2130, MTI-GlobalStem) to the DNA+Opti-MEM mix from above. The mix was incubated for 10 minutes at room temperature before distributing it to the cells. The next day the cells were fed with fresh media. At ~40 hours after transfection, the cells were selected with 0.5-1.0 μg/mL of puromycin for 24 hours. Following selection, the cells were allowed to recover for 5-7 days and then passaged as 500-1000 single cells per well of a 6 well plate for colony formation. These cells were maintained for 7-10 days before colony picking and PCR analysis.

#### Mouse stem cells

mESCs were passaged and plated as 100K per well of a 6 well plate the day before transfection. The following day, (for one well) 1 μg of all-in-one Cas9 plasmid and 2 μg of donor plasmid were combined with Opti-MEM to a total volume of 145 μL. After mixing, 5 μL of DNA-In Stem were added to the solution for a 15 minute incubation at room temperature before distribution to a plate well. 24 hours after transfection the cells were selected with 3 μg/mL of puromycin for 24 hours. Following selection, the cells were allowed to recover for 1 day and then passaged as 750 single cells per well of a 6 well plate for colony formation. These cells were maintained for 5 days before colony picking and PCR analysis

Note MTI-GlobalStem is now part of Thermo Fisher Scientific which recommends that DNA-In Stem should be replaced by Lipofectamine Stem (STEM00015).

### Flow cytometry

The gates for reporter negative cells were set based upon values of untransfected PSCs of the line being analyzed. Side scatter height versus width linear alignment filters were used to minimize cell aggregates. Cells were dissociated to a single cell suspension using Accumax (07921, Stemcell Technologies). The SH-800 Cell Sorter (Sony Biotechnology, San Jose, CA, https://www.sonybiotechnology.com) was used for analysis.

### Microscopy

Fluorescence images were taken using the Eclipse TE-2000S inverted microscope (Nikon, Tokyo, Japan, http://www.nikon.com) or the EVOS FL Auto 2 Cell Imaging System (Thermo Fisher Scientific).

### Karyotyping/off-target effects

Karyotyping analysis of the EP1-RGC reporter line was performed using the KaryoStat™ Analysis Service (Thermo Fisher Scientific, A38153) and no chromosomal aberrations were found. To test the EP1-RGC reporter line for CRISPR off-target effects, we performed PCR and sequenced the 4 most likely off-targets as assessed by the CCTop online tool(44) (https://crispr.cos.uni-heidelberg.de).

### Stem cell differentiation

#### Human

A homozygous *BRN3B-P2A-tdTomato-P2A-Thy1.2* reporter clone was isolated from EP1 hiPSCs and differentiated to RGCs using our previously published protocol(11).

#### Mouse

A homozygous *Six6-P2A-eGFP* reporter clone was isolated from E14TG2a mESCs and differentiated to optic vesicles using the published protocol(33).

### Statistical analysis

An unpaired two tailed t-test was used as indicated. Statistics were analyzed in GraphPad Prism7.

## Acknowledgments

This work was supported by grants from the NIH (including 5P30EY001765), Maryland Stem Cell Research Fund, Foundation Fighting Blindness, BrightFocus Foundation, Thome Foundation, and unrestricted funds from Research to Prevent Blindness, Inc., and generous gifts from the Guerrieri Family Foundation.

---

**S1 Fig. EP1-RGC reporter line off-target mutation and karyotype analysis.**

(A) The four most likely *in-silico* predicted off-target mutations for the *BRN3B* gRNA were sequenced and confirmed to be WT. (B) Karyotyping analysis using KaryoStat^TM^ found no chromosomal aberrations.

**Author contributions**
V.M.S., X.C., and D.J.Z. conceived the study. V.M.S. and X.C. contributed to reagent generation, data acquisition and interpretation, and manuscript preparation. C.W. generated the EP1 hiPSC RGC reporter line. Y.D. contributed to reagent generation. V.M.S. wrote the manuscript with contributions from all other authors.

## References

1. Sluch VM, Zack DJ. Stem cells, retinal ganglion cells and glaucoma. Dev Ophthalmol. 2014;53:111–21.

2. Chamling X, Sluch VM, Zack DJ. The Potential of Human Stem Cells for the Study and Treatment of Glaucoma. Invest Ophthalmol Vis Sci. 2016;57(5):ORSFi1–6.

3. Shi Y, Inoue H, Wu JC, Yamanaka S. Induced pluripotent stem cell technology: a decade of progress. Nat Rev Drug Discov. 2017;16(2):115–30.

4. Schwartz SD, Regillo CD, Lam BL, Eliott D, Rosenfeld PJ, Gregori NZ, et al. Human embryonic stem cell-derived retinal pigment epithelium in patients with age-related macular degeneration and Stargardt’s macular dystrophy: follow-up of two open-label phase 1/2 studies. Lancet. 2015;385(9967):509–16.

5. Mandai M, Watanabe A, Kurimoto Y, Hirami Y, Morinaga C, Daimon T, et al. Autologous Induced Stem-Cell-Derived Retinal Cells for Macular Degeneration. N Engl J Med. 2017;376(11):1038–46.

6. da Cruz L, Fynes K, Georgiadis O, Kerby J, Luo YH, Ahmado A, et al. Phase 1 clinical study of an embryonic stem cell-derived retinal pigment epithelium patch in age-related macular degeneration. Nat Biotechnol. 2018;36(4):328–37.

7. Kashani AH, Lebkowski JS, Rahhal FM, Avery RL, Salehi-Had H, Dang W, et al. A bioengineered retinal pigment epithelial monolayer for advanced, dry age-related macular degeneration. Sci Transl Med. 2018;10(435).

8. Gaj T, Gersbach CA, Barbas CF, 3rd. ZFN, TALEN, and CRISPR/Cas-based methods for genome engineering. Trends Biotechnol. 2013;31(7):397–405.

9. Soldner F, Stelzer Y, Shivalila CS, Abraham BJ, Latourelle JC, Barrasa MI, et al. Parkinson-associated risk variant in distal enhancer of alpha-synuclein modulates target gene expression. Nature. 2016;533(7601):95–9.

10. Phillips MJ, Capowski EE, Petersen A, Jansen AD, Barlow K, Edwards KL, et al. Generation of a rod-specific NRL reporter line in human pluripotent stem cells. Sci Rep. 2018;8(1):2370.

11. Sluch VM, Chamling X, Liu MM, Berlinicke CA, Cheng J, Mitchell KL, et al. Enhanced Stem Cell Differentiation and Immunopurification of Genome Engineered Human Retinal Ganglion Cells. Stem Cells Transl Med. 2017;6(11):1972–86.

12. Sluch VM, Davis CH, Ranganathan V, Kerr JM, Krick K, Martin R, et al. Differentiation of human ESCs to retinal ganglion cells using a CRISPR engineered reporter cell line. Sci Rep. 2015;5:16595.

13. Maruotti J, Sripathi SR, Bharti K, Fuller J, Wahlin KJ, Ranganathan V, et al. Small-molecule-directed, efficient generation of retinal pigment epithelium from human pluripotent stem cells. Proc Natl Acad Sci U S A. 2015;112(35):10950–5.

14. Yang L, Guell M, Byrne S, Yang JL, De Los Angeles A, Mali P, et al. Optimization of scarless human stem cell genome editing. Nucleic Acids Res. 2013;41(19):9049–61.

15. Zhu Z, Verma N, Gonzalez F, Shi ZD, Huangfu D. A CRISPR/Cas-Mediated Selection-free Knockin Strategy in Human Embryonic Stem Cells. Stem Cell Reports. 2015;4(6):1103–11.

16. Ding Q, Regan SN, Xia Y, Oostrom LA, Cowan CA, Musunuru K. Enhanced efficiency of human pluripotent stem cell genome editing through replacing TALENs with CRISPRs. Cell Stem Cell. 2013;12(4):393–4.

17. Ding Q, Lee YK, Schaefer EA, Peters DT, Veres A, Kim K, et al. A TALEN genome-editing system for generating human stem cell-based disease models. Cell Stem Cell. 2013;12(2):238–51.

18. Yu C, Liu Y, Ma T, Liu K, Xu S, Zhang Y, et al. Small molecules enhance CRISPR genome editing in pluripotent stem cells. Cell Stem Cell. 2015;16(2):142–7.

19. Lin S, Staahl BT, Alla RK, Doudna JA. Enhanced homology-directed human genome engineering by controlled timing of CRISPR/Cas9 delivery. Elife. 2014;3:e04766.

20. He X, Tan C, Wang F, Wang Y, Zhou R, Cui D, et al. Knock-in of large reporter genes in human cells via CRISPR/Cas9-induced homology-dependent and independent DNA repair. Nucleic Acids Res. 2016;44(9):e85.

21. Gonzalez F, Zhu Z, Shi ZD, Lelli K, Verma N, Li QV, et al. An iCRISPR platform for rapid, multiplexable, and inducible genome editing in human pluripotent stem cells. Cell Stem Cell. 2014;15(2):215–26.

22. Kim S, Kim D, Cho SW, Kim J, Kim JS. Highly efficient RNA-guided genome editing in human cells via delivery of purified Cas9 ribonucleoproteins. Genome Res. 2014;24(6):1012–9.

23. Petris G, Casini A, Montagna C, Lorenzin F, Prandi D, Romanel A, et al. Hit and go CAS9 delivered through a lentiviral based self-limiting circuit. Nat Commun. 2017;8:15334.

24. Chen Y, Cao J, Xiong M, Petersen AJ, Dong Y, Tao Y, et al. Engineering Human Stem Cell Lines with Inducible Gene Knockout using CRISPR/Cas9. Cell Stem Cell. 2015;17(2):233–44.

25. Merkle FT, Neuhausser WM, Santos D, Valen E, Gagnon JA, Maas K, et al. Efficient CRISPR-Cas9-mediated generation of knockin human pluripotent stem cells lacking undesired mutations at the targeted locus. Cell Rep. 2015;11(6):875–83.

26. Meier ID, Bernreuther C, Tilling T, Neidhardt J, Wong YW, Schulze C, et al. Short DNA sequences inserted for gene targeting can accidentally interfere with off-target gene expression. FASEB J. 2010;24(6):1714–24.

27. Wang G, Yang L, Grishin D, Rios X, Ye LY, Hu Y, et al. Efficient, footprint-free human iPSC genome editing by consolidation of Cas9/CRISPR and piggyBac technologies. Nat Protoc. 2017;12(1):88–103.

28. Bhise NS, Wahlin KJ, Zack DJ, Green JJ. Evaluating the potential of poly(beta-amino ester) nanoparticles for reprogramming human fibroblasts to become induced pluripotent stem cells. Int J Nanomedicine. 2013;8:4641–58.

29. Wahlin KJ, Maruotti JA, Sripathi SR, Ball J, Angueyra JM, Kim C, et al. Photoreceptor Outer Segment-like Structures in Long-Term 3D Retinas from Human Pluripotent Stem Cells. Sci Rep. 2017;7(1):766.

30. Mitzelfelt KA, McDermott-Roe C, Grzybowski MN, Marquez M, Kuo CT, Riedel M, et al. Efficient Precision Genome Editing in iPSCs via Genetic Co-targeting with Selection. Stem Cell Reports. 2017;8(3):491–9.

31. Shy BR, MacDougall MS, Clarke R, Merrill BJ. Co-incident insertion enables high efficiency genome engineering in mouse embryonic stem cells. Nucleic Acids Res. 2016;44(16):7997–8010.

32. Volkner M, Zschatzsch M, Rostovskaya M, Overall RW, Busskamp V, Anastassiadis K, et al. Retinal Organoids from Pluripotent Stem Cells Efficiently Recapitulate Retinogenesis. Stem Cell Reports. 2016;6(4):525–38.

33. Assawachananont J, Mandai M, Okamoto S, Yamada C, Eiraku M, Yonemura S, et al. Transplantation of embryonic and induced pluripotent stem cell-derived 3D retinal sheets into retinal degenerative mice. Stem Cell Reports. 2014;2(5):662–74.

34. Roy A, de Melo J, Chaturvedi D, Thein T, Cabrera-Socorro A, Houart C, et al. LHX2 is necessary for the maintenance of optic identity and for the progression of optic morphogenesis. J Neurosci. 2013;33(16):6877–84.

35. Tetreault N, Champagne MP, Bernier G. The LIM homeobox transcription factor Lhx2 is required to specify the retina field and synergistically cooperates with Pax6 for Six6 trans-activation. Dev Biol. 2009;327(2):541–50.

36. Nakatake Y, Fujii S, Masui S, Sugimoto T, Torikai-Nishikawa S, Adachi K, et al. Kinetics of drug selection systems in mouse embryonic stem cells. BMC Biotechnol. 2013;13:64.

37. Oji A, Noda T, Fujihara Y, Miyata H, Kim YJ, Muto M, et al. CRISPR/Cas9 mediated genome editing in ES cells and its application for chimeric analysis in mice. Sci Rep. 2016;6:31666.

38. Den Hartogh SC, Schreurs C, Monshouwer-Kloots JJ, Davis RP, Elliott DA, Mummery CL, et al. Dual reporter MESP1 mCherry/w-NKX2-5 eGFP/w hESCs enable studying early human cardiac differentiation. Stem Cells. 2015;33(1):56–67.

39. Steyer B, Bu Q, Cory E, Jiang K, Duong S, Sinha D, et al. Scarless Genome Editing of Human Pluripotent Stem Cells via Transient Puromycin Selection. Stem Cell Reports. 2018;10(2):642–54.

40. Li K, Wang G, Andersen T, Zhou P, Pu WT. Optimization of genome engineering approaches with the CRISPR/Cas9 system. PLoS One. 2014;9(8):e105779.

41. Guo Q, Mintier G, Ma-Edmonds M, Storton D, Wang X, Xiao X, et al. ‘Cold shock’ increases the frequency of homology directed repair gene editing in induced pluripotent stem cells. Sci Rep. 2018;8(1):2080.

42. Zhang JP, Li XL, Li GH, Chen W, Arakaki C, Botimer GD, et al. Efficient precise knockin with a double cut HDR donor after CRISPR/Cas9-mediated double-stranded DNA cleavage. Genome Biol. 2017;18(1):35.

43. Naito Y, Hino K, Bono H, Ui-Tei K. CRISPRdirect: software for designing CRISPR/Cas guide RNA with reduced off-target sites. Bioinformatics. 2015;31(7):1120–3.

44. Stemmer M, Thumberger T, Del Sol Keyer M, Wittbrodt J, Mateo JL. CCTop: An Intuitive, Flexible and Reliable CRISPR/Cas9 Target Prediction Tool. PLoS One. 2015;10(4):e0124633.

